# iHyd-ProSite: A novel Computational Approach for Identifying Hydroxylation Sites in Proline Via Mathematical Modeling

**DOI:** 10.1101/2020.03.03.974717

**Authors:** Muhammad Khalid Mahmood, Asma Ehsan, Yaser Daanial Khan

## Abstract

In various cellular functions, post translational modifications (PTM) of protein play a vital role. The addition of certain functional group through a covalent bond to the protein induces PTM. The number of PTMs are identified which are closely linked with diseases for example cancer and neurological disorder. Hydroxylation is one of the PTM, modified proline residue within a polypeptide sequence. The defective hydroxylation of proline causes absences of ascorbic acid in human which produce scurvy, and many other dominant health issues. Undoubtedly, the prediction of hydroxylation sites in proline residues is of challenging frontier. The experimental identification of hydroxyproline site is quite difficult, high-priced and time-consuming. The diversity in protein sequences instigates to develop a computational tool to identify hydroxylated site within short time with excellent prediction accuracy to handle such proteomics problems. In this work a novel in silico predictor is developed through rigorous mathematical modeling to identify which site of proline is hydroxylated and which site is not? Then performance of the predictor was verified using three validations tests, namely self-consistency test, cross-validation test and jackknife test over the benchmark dataset. A comparison was established for jackknife test with the previous methods. In comparison with previous predictors the proposed tool is more accurate than the existing techniques. Hence this scheme is highly useful and inspiring in contrast to all previous predictors.

## Introduction

In mammals collagens are extremely abundant protein comprised of proline modified residue during the chemical process such as hydroxylation and produce hydroxyproline [1]. Collagens are stringy and long in nature, most of the protein in mammals consists of almost a quarter part of collagen [2]. In the treatment of wound healing [3], burn and cosmetic surgeries [4, 5] collagen mainly works as a medicinal drug. Most of the dominant human diseases like stomach and lung cancer [6, 7] are closely related with the defects and irregularities in hydroxylation process. Thus the identification of hydroxyproline (HyP) sites in proteins gives valuable data helpful to both biomedical research and drug development [8]. Hydroxyproline obtained by the addition of a hydroxyl group (-OH) to the proline (P) residue modifies the CH group into the COH group [8] (see Fig. 1).

**Fig 1.**
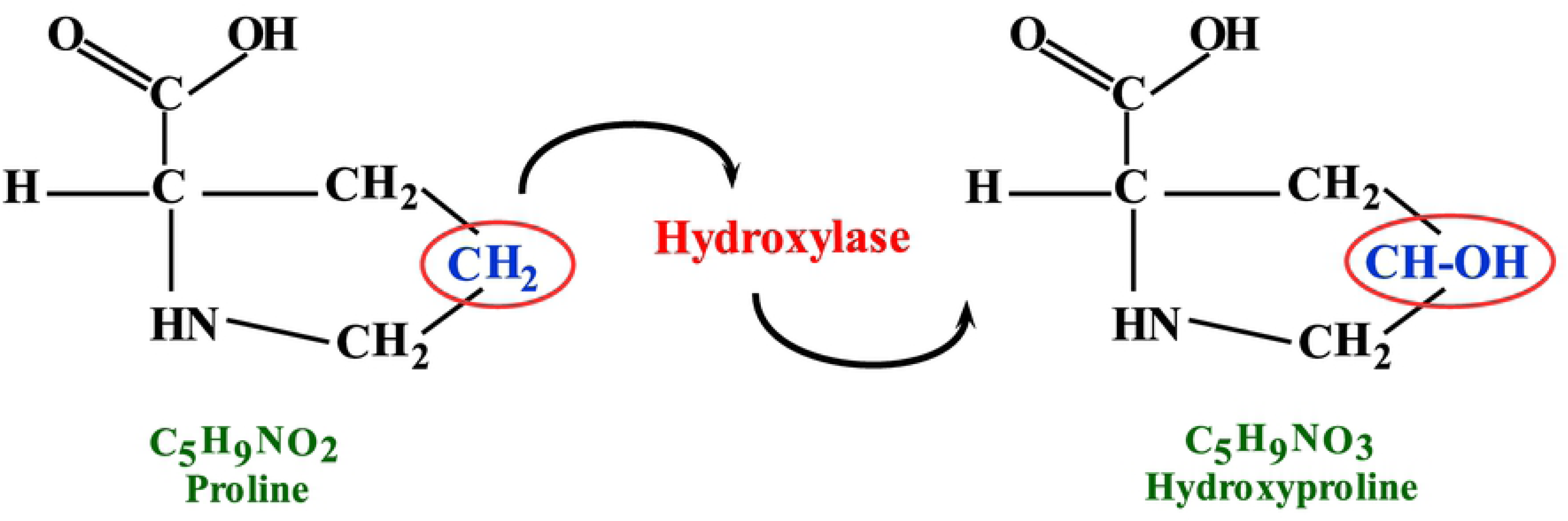
Conversion of proline (Pro) residue into hydroxyproline. The figure is to show that, hydroxylation action attaches the -OH group to proline (Pro) to convert CH group to COH and modify proline residue into hydroxyproline.

The number of scientists has been making their contributions [9–12] in order to understand the cellular biological process and finding out medicines for cancer and for various other diseases. The prediction of hydroxylation sites in the lab by using method mass spectrometry [13] is difficult to conduct, expensive and very lengthy process. Every day, the large number of protein sequences is collected in the data bank and to classify them according to their functional properties is a crucial. It is highly worthy to build an efficient computational predictor for the classification of targeted hydroxylation sites within polypeptide sequences with improved prediction accuracy. Many researchers have been developed a couple of methods in this regard. Still, all these previous methods are insufficient to incorporate all components of features vague in the polypeptide sequence that become difficult to get exact prediction. Many scientists had been shown their great interest in hydroxylation process. Colgrave, et al. [1] was computed quantification of hydroxyproline by using multiple reaction monitoring mass spectrometry. In order to understand the microbial activity and their communities, a mathematical model has been developed [14]. A system was defined to study the insufficiency of collagen in connective tissues that encountered by lack of ascorbic acid [15].

Halme et al. [16] and kiviriko et al. [17] were explained the separation and classification of extremely purified protocollagen proline hydroxylase as well as proline hydroxylation in synthetical proteins with pure procollagen hydroxylase. In human proteome, the functional character of proline and polyproline based on distribution, frequency and positioning was investigated by Morgan, et al. [18]. Yamauchi et al. [19] were elaborated the Hydroxylation of lysine and cross-linking of collagens. By using a position weight of 8 high-quality amino acid indices and via support vector machines, Shi, Shao-Ping, et al. [20] were proposed a novel technique named as PredHydroxy for the forecast of the proline and lysine hydroxylation locales. Moreover, the functional study of proline with mutable surroundings and the metabolism of proline, hydroxyproline were examined in [21, 22]. ZR Yang [23] developed a tool for the prediction of hydroxyproline sites by utilizing support vector machine. A sequence-based formulation for identifying hydroxyproline and hydroxylysine were developed by Hu, Le-Le, et al. [24]. Using dipeptide position and specific propensity into pseudo amino acid composition Xu, Yan, et al. [8] predicted hydroxyproline and hydroxylysine in proteins. Qiu, Wang-Ren, et al. [25] was suggested an enhanced method over this technique by assimilating a sequence coupled effect into general PseAAC.

## Materials and methods

### Benchmark Dataset

The acquiring of benchmark dataset is critical, as indicated by Chou’s 5-step rule [26] that prompts the attaining of a powerful, assorted and improved dataset. In order to obtain a stringent benchmark dataset, two resources have been used in the current study. One of the supported datasets is obtained from the universal protein database http://www.uniprot.org/, while the other dataset is borrowed from a posttranslational modification database dbPTM 3.0 [42]. Thus, a stringent benchmark datasets are obtained by employing the following two steps.

**Step-1**: The extracted dataset from UniProt database, contains two sets of protein sequences. One of the set represents hydroxylated protein sequences at proline site and labeled as positive sample. Likewise, other set consists of non-hydroxylated protein sequences at proline site, tagged as negative sample. An inquiry is produced to choose polypeptide sequences in the PTM/processing field as hydroxyproline. Records construed with any experimental assertion in Feature Table (FT) were only chosen. After a thorough selection of the described query, a stringent benchmark dataset of hydroxyproline was obtained. There were found records of 816 and 24980 for hydroxylated and non-hydroxylated sequences. The records were reduced to 782 and 24971 respectively, after removing duplicates.

**Step-2**: Likewise, to obtain another stringent benchmark dataset the dbdtm 3.047 were utilized. The dataset was effectively accessible in FASTA format and advantageously were downloaded for hydroxylation (hydroxylated and non-hydroxylated). There were discovered 226 hydroxylated records and 3,865 non-hydroxylated samples. The primary dataset of hydroxylated and non-hydroxylated proline sites can be found in Supplementary Tables S1, S2, S3 and S4 separately.

### Method

In order to identify target proline sites with hydroxylation, an excellent methodology is proposed as indicated in the Chou’s second and third step [26]. This technique is developed by incorporating all indispensible components of polypeptide sequences that can perfectly indicate their correlation to assemble the sequence in an effective way. The alternate formulation was also employed by Ehsan et al. [31, 32], impart as prominent prediction rate in proteomics problems. Consider a protein sample ℂ consists of Z amino acid residues.

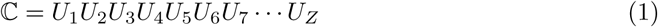

Where *U*_1_ indicates the first amino acid residue with in string ℂ, *U*_2_ is the second amino acid component, *U*_3_ is third component and so on up till *U*_*Z*_ last amino acid residue of the protein sequence ℂ as given in (1). In this methodology the formulation is handled by upholding sequence information. By considering the sequence order effect and composition of each term of sequence an advantageous factor introduced, known as weight factor. This is used to maintain position and composition of all components of sequence and denoted by *T*_*i*_. While the significant terms L, M, and N indicate the count factor of each residue with its contiguous residues in both forward and reverse direction. The weight factor *T*_*i*_ is characterized as: the product of the position with the occurrence of the each term of the sequence among the similar residues. This whole scheme is based on expression (2) to handle the diverse length of the polypeptide sequences. The term {*L* + *M* + *N*} describes the weighted mean of all possible coupling between similar residues that is after the first residue and before it occurred again.

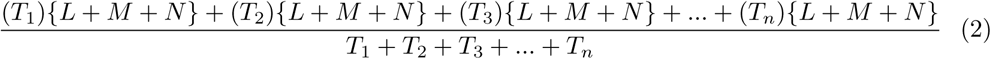

Where the weight factors *T*_1_, *T*_2_, *T*_3_, …, *T*_*n*_ depends upon the repeated terms of the sequence of type 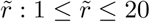. Each *T*_*i*_ is estimated between the two consecutive terms *T*_*i*_ and *T*_*i*+1_ before 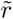 and when it occurred again. All weight factors characterization of amino acid of type 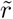 in terms of mathematical form is represented as: 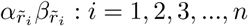 (*α* represent the occurrence of residue of type 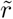 at their corresponding positions *β* in the sequence varies with i, represents n time appearance of 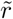) and L, M, N is the estimated count of the three correlated factors in both forward and backward direction from 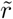 with its contiguous residues except 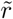 until 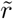 appeared again. The demonstration of the above process is mathematically expanded in Eqs. (3) and (4) which collectively set up a mean, 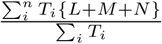 as given in *T* expression (2). Whereas *i* depends upon the number of compositions of residue of type 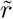 in concatenation. Moreover, non-occurrence will assign zero value corresponding to the weight factor, so this weight is neglected and only considered the weight factors for has occurred objects.

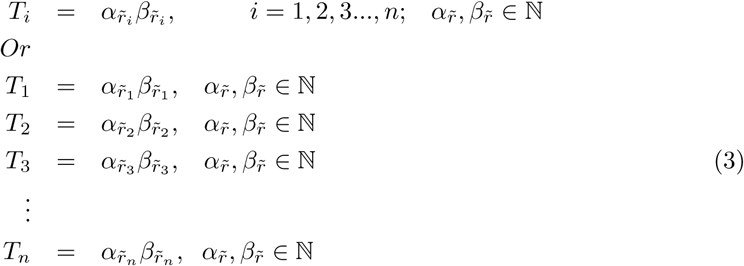

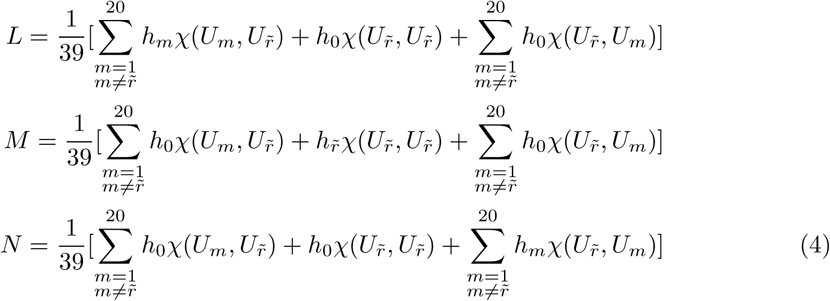

Where *h*_*m*_, 1 ≤ *m* ≤ 20 symbolizes the repeated coupling function *χ* of residue 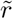 with all amino acid residues also 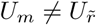. For non-occurrence, it is denoted by *h*_0_. The coupling function of 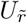 with itself is denoted by 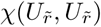 and the frequency of this pair is represented by 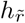. For the sake of convenience, consider *χ*(*U*_*i*_, *U*_*j*_) = *ω*_*i,j*_; *i* = *j* = 1, 2, 3, …, 20. Whereas *ω*_*i,j*_ represents all possible coupling factors for all amino acid residue with each other. A complete interpretation for all possible correlation is given in matrix representation (5) and in term of L, M and N separately assigned in (6). The matrix (6) is adopted by constraint (7), when the pair *χ*(*U*_*i*_, *U*_*j*_) appeared, then *ω*_*i,j*_ gives 1 otherwise it is attributed as number zero.

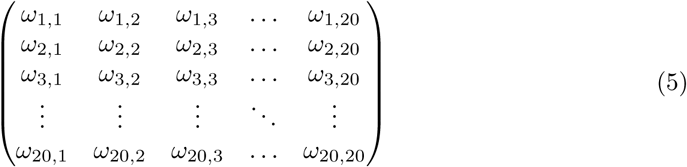

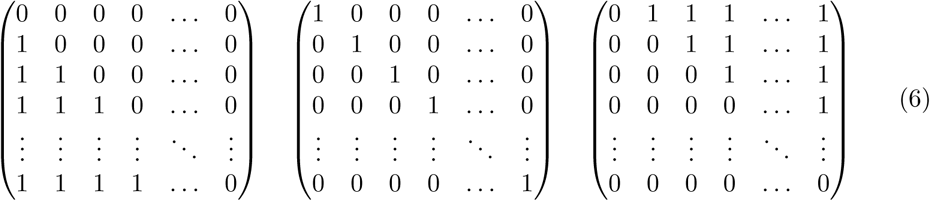

Where

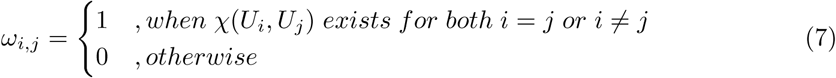

Use Eqs. (3) and (4) in expression (2) to get feature vector reflecting the residue 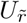 of type 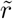. The desired feature vector is given in Eq. (8) and compactly defined in Eq. (9).

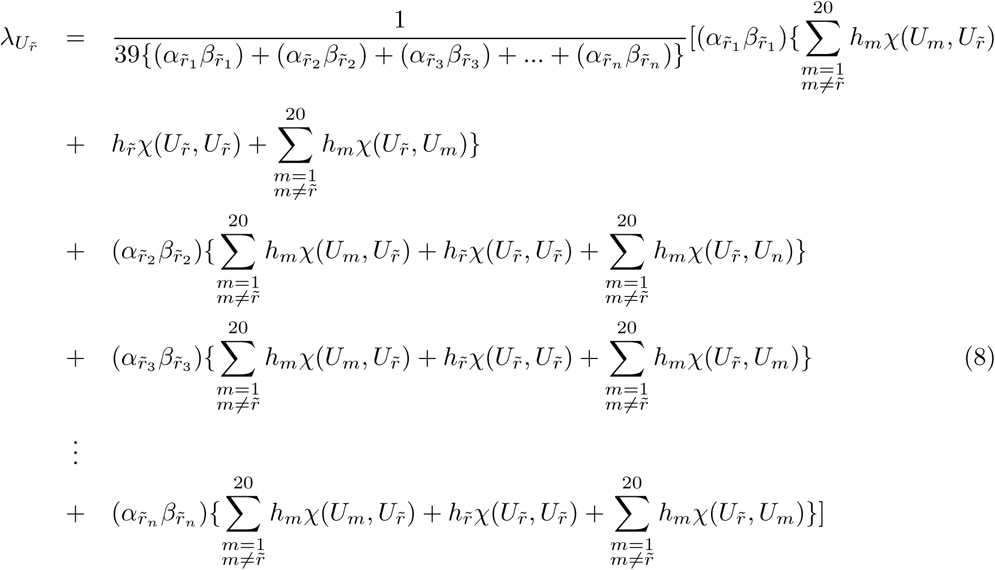

Or

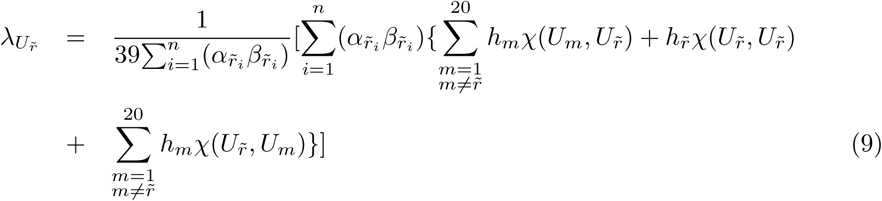

In order to understand the mechanism of proposed model consider ith term *U*_*i*_ of sequence (1), reflects the first alphabetical letter of amino acid residues, say, “A”. Notice its occurrences as well as its corresponding positions in the sequence. *U*_*i*_ makes pairing with its contiguous residues in reverse and forward direction. The *ith* residue in term of *χ*(*U*_*m*_, *U*_*i*_) and *χ*(*U*_*i*_, *U*_*m*_) represented by green and blue curved lines (see Fig. 2). This process will be continued until next *U*_*j*_ occurs at *jth* position such that *U*_*i*_ = *U*_*j*_ = *A*. Similarly, the same steps will be conducted for *U*_*j*_. The feature component corresponding to residue “A” is interpreted in Eq. (10).

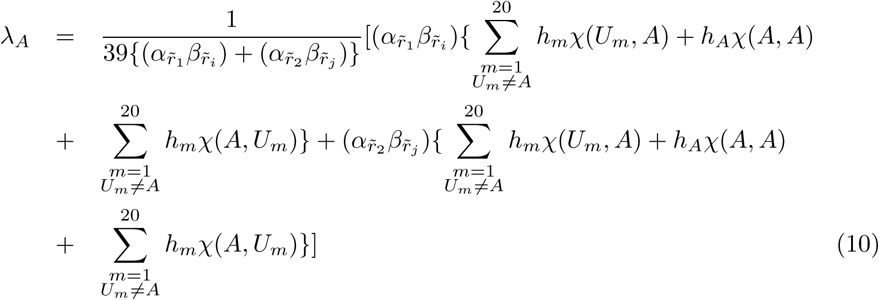

**Fig 2.**
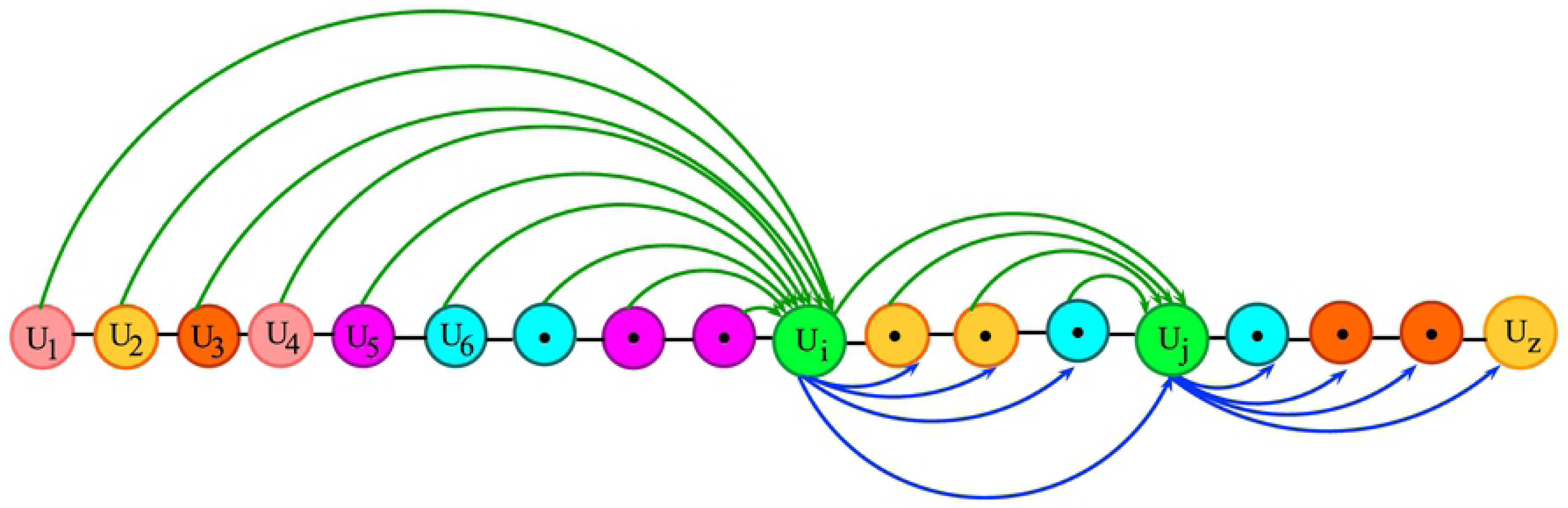
A mechanism for sequence formulation. The figure is to show the graphical demonstration of scheme feature vector for the residue “A”, representing how “A” make pairs with its contiguous residues in both directions up to next residue.

Where the numeral values *m* = 1, 2, 3…, 20 indicates the twenty amino acid residues of alphabetical order. For more convenience, assume that *U*_1_, *U*_2_, *U*_3_, …, *U*_20_ represents 20 amino acids of alphabetical order labeled as: A, C, D, E, F, G, H, I, K, L, M, N, P, Q, R, S, T, V, W and Y and *U*_21_ onwards the 20 residues cyclically repeats themselves then *λ*_1_,*λ*_2_,*λ*_3_,…,*λ*_20_ be their corresponding feature components. The set of twenty feature components is given in Eq. (11) as follows.

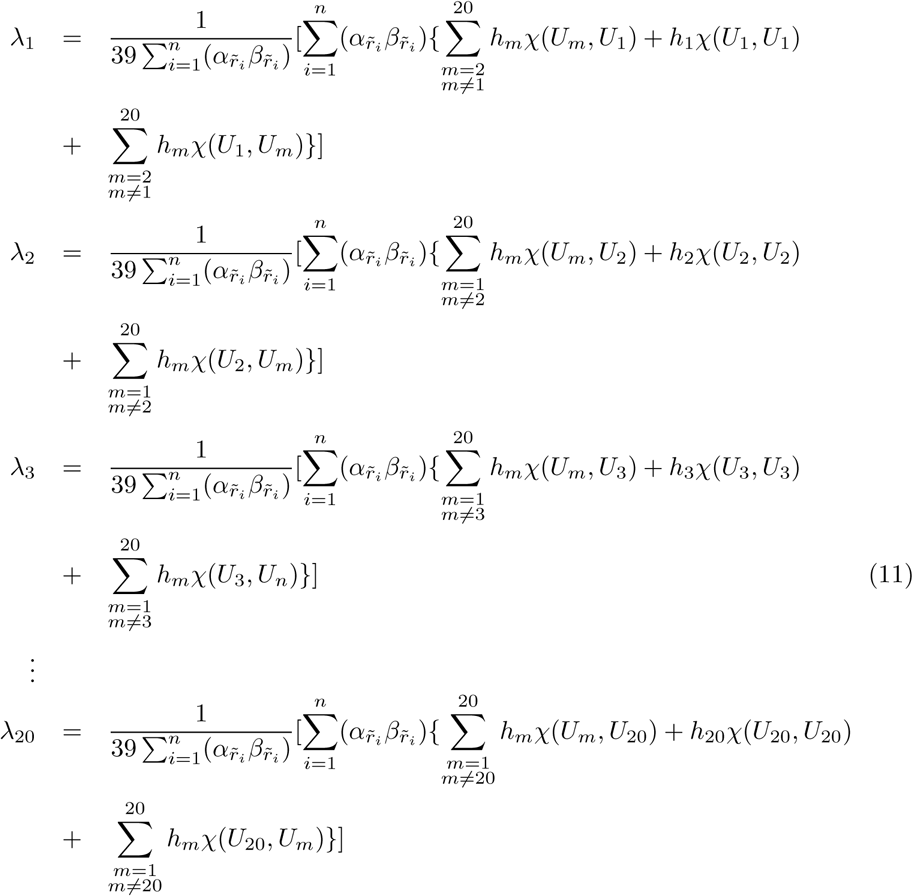

The above set of twenty feature vectors depends upon three properties of amino acids such that, hydrophobicity, hydrophilicity and side chain mass of amino acids, can be calculated by employing Eqs. (12) to (14). These equations can expand as per choice of attributes of amino acids other than these three properties of amino acid. For extended properties *l* of amino acids a compact representation is elaborated in Eq. (15).

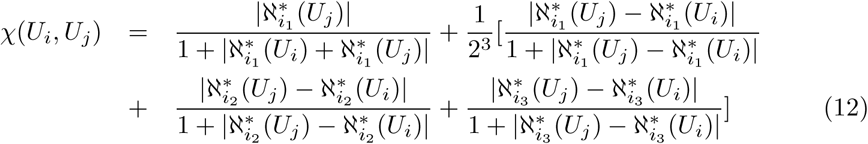

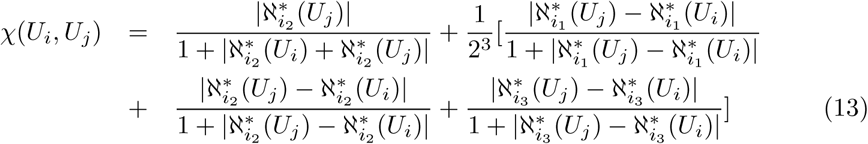

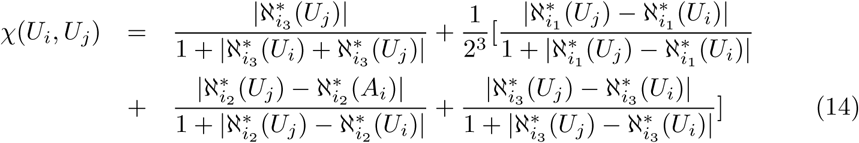

Or

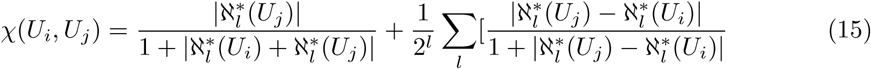

Whereas 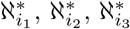 are the values of hydrophobicity, hydrophilicity and side-chain mass of amino acid residues that are normalized by using Eq. (16) against the pair of *U*_*i*_ and *U*_*j*_. The normalization index that is used to normalize the values given in Eqs. (12) to (14) lies between (-S, S), where S is the normalizing count for 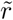 amino acids. Here the number 5 is used for normalization. The original values for hydrophobicity and hydrophilicity were taken from the main source, employed by Ehsan at el. [31, 32], while the values for side-chain mass of amino acid residues was taken from any text book of biochemistry.

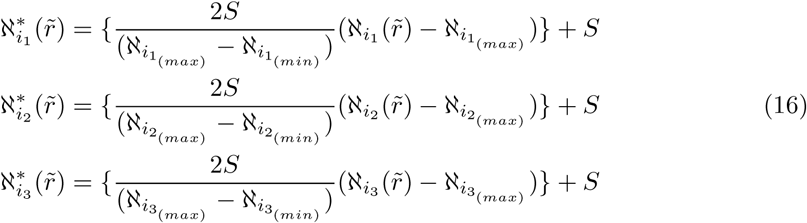

The provided modelling consist of 100 dimensions by comprising the protein features and to classify them according to their functional properties and attributes. These features are divided as: the very first twenty feature vectors corresponds to hydrophobic property, while next twenty matches for hydrophilic attribute. Similarly the succeeding twenty vectors indicate side chain mass property of amino acid residues and last forty vectors related to the position and composition of each amino acids. For the identification of diverse protein sequence this novel technique establishes a wonderful result. For the sake of classification these extracted feature vectors are further passed through a training-testing process by using the rigorous classifier, neural networks (NN).

The neural network is an extraordinary tool for decision making problems and to classify patterns in available diversified data sets. It is typically arranged in layers and learn from its experience using input data and able to modify their weights according to provided data. Subsequent to the training process is finished the system apparently acts such that makes it fit to arrange each given input inside a worthy level of precision. Its connectionist structural design comprises of 100 input layer neurons, 50 hidden layer neurons and two output neurons that classify hydroxylated and non-hydroxylated protein samples. The back propagation method was used for training of the multilayered neural network. In order to get the higher prediction rate and to decrease the error rate a gradient descent method was employed with adaptive learning rate.

The results were simulated on MATLAB R2017 version and were duplicated on python ver 3.6 platform along with Scikit Learn 0.20 for neural network training and simulation bearing identical results. This procedure is done in the flowchart as given underneath (see Fig. 3).

**Fig 3.**
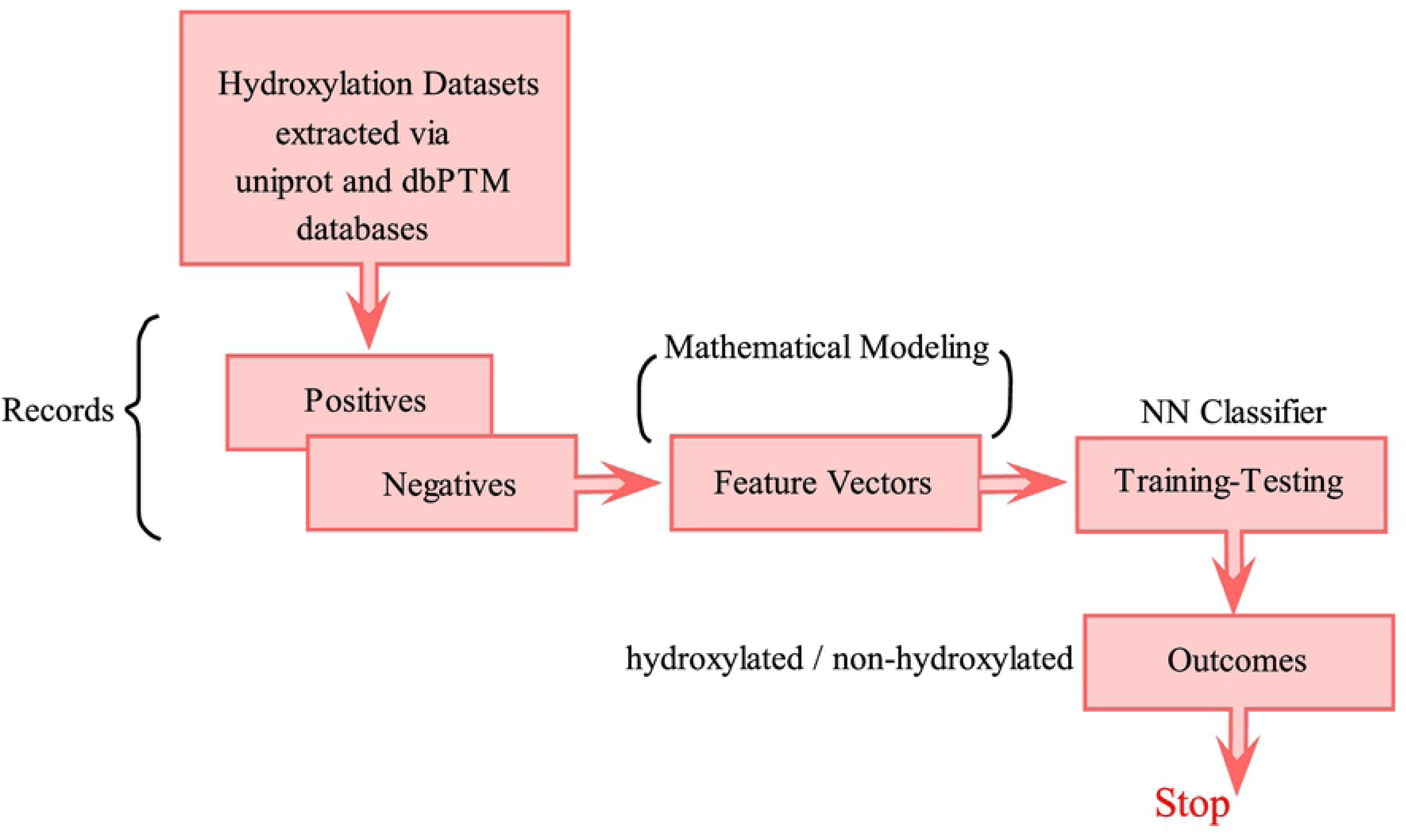
Flowchart describes the training and validation process. The prediction process done in the following steps (a) extraction of two class dataset (b) extract feature vectors using the proposed tool (c) train/test dataset using neural network classifier.

## Results

In order to build up a beneficial predictor for an organic development, the Chou’s 5-step rule [26] are noticeable. Undoubtedly, it is useful to develop a new predictor by employing Chou’s 5-step rule. A number of researchers [27–30] had used this method in their work, published very recently. The prediction analysis is done in some steps: firstly the stringent benchmark data set is collected for training and testing purpose of proposed predictor, in a second step, a powerful mathematical tool developed that select the major and most significant features of the amino acid polymers. Then the developed feature vector incorporated into an identifying formulation for the sake of training. When the process of training is completed, the trained model is completely tested and validated. Finally, a web-server is created for open use of the proposed predictor. In the current study, the initial four steps have been carefully performed, while, the last step has been kept open for future work.

### Statistical Measures

To evaluate the performance of the proposed model “iHyd-ProSite”, a set of four metrics are followed, which were employed by Ehsan et al. [31,32]. The following these four metrics are: sensitivity (Sn), specificity (Sp), accuracy (Acc), and Matthews correlation coefficients (MCC) respectively, used for proposed algorithm evaluation. Either the set of traditional metrics copied from maths books or the intuitive metrics derived from the Chou’s symbols [33, 34] is valid only for the single-label systems (where each sample only belongs to one class). For the multilabel systems (where a sample may simultaneously belong to several classes), whose existence has become more frequent in system biology [28, 35], system medicine [36] and biomedicine [37], a completely different set of metrics as defined in the study represented as a reference [38] is absolutely needed.

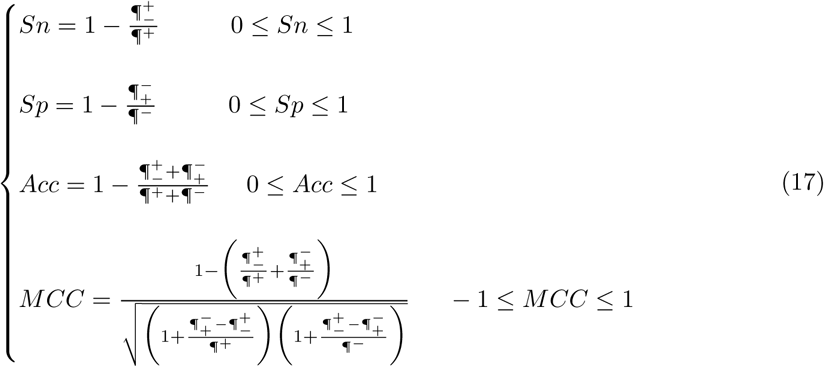

Consider ¶^+^ and ¶^−^ represents the all correctly predicted positive and negative records for hydroxylated and non-hydroxylated site in proline (Pro) within polypeptide sequences. Similarly, if the positive records are wrongly predicted as negative records, is denoted by 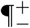 and when negative records are wrongly predicted in terms of positive records is represented by 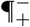. It is relevant to discuss the following cases of the above Eq. (17). If ¶^+^_−_ = 0, then there is no incorrectly predicted positive records as negative records, then produce *Sn* = 1. In another case when ¶^+^_−_ = ¶^+^ indicating that all positive records were incorrectly predicted in terms of negative records so, the sensitivity is *Sn* = 0. Similarly for ¶^−^_+_ = 0 gives specificity *Sp* = 1 representing no any negative record peptide was incorrectly predicted in terms of positive records, while for ¶^−^_+_ = ¶^−^ gives specificity *Sp* = 0 denoting all negative instances were wrongly predicted as positive instances. On the other hand the prediction accuracy *Acc* = 1 when, there is no any incorrectly predicted sequences were found both positive and negative cases such that, ¶^+^_−_ = ¶^−^_+_ = 0. When ¶^+^_−_ = ¶^+^ and ¶^−^_+_ = ¶^−^, brings out misclassification so the overall accuracy is *Acc* = 0. Furthermore, the performance of binary classifications is often measured by Matthew correlative coefficient (MCC). There are three cases here, for ¶^+^_−_ = ¶^−^_+_ = 0 indicating that there is no incorrectly predicted record were found for both positive and negative instances so we obtain *MCC* = 1. In the second case when 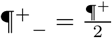 and 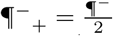 we obtain *MCC* = 0 indicating the inaccurate prediction. Lastly, when ¶^+^_−_ = ¶^+^ and ¶^−^_+_ = ¶^−^ we obtain *MCC* = −1 denoting the totally wrong binary classification and disagreement between observed and predicted values.

### Renowned Validation Tests

In order to validate the quality of the proposed predictor the following three test methods, self-consistency test, K-fold cross validation test and jackknife test are often used. These tests are applied to score the metrics given in Eq. (17). To approve the predictor’s quality these tests are viewed as valuable. By employing the statistical measures, a comparison was made using the jackknife test with the existing predictors [8, 25]. In the current study, to test the performance of proposed scheme all above test methods were employed. Additionally, for validation purpose the benchmark datasets were taken from two sources, one is from uniprot and other one is from dbptm. The results obtained by using both datasets are given in Table 1. Table 1 is divided into two main columns. The first column is explaining the values of metrics for dbptm dataset by employing all above tests. Whereas, second column is giving the values in validation of uniprot dataset. It is noticed that the values for MCC were 0.91 and 0.90 for jackknife test. The comparison of the proposed scheme with the existing techniques for jackknife test is given below.

**Table 1.** The values Table by using proposed predictor “iHyd-ProSite”. The results obtained by employing the proposed predictor by using self-consistency test, cross-validation, test and jackknife test on the set of metrics for dbptm and uniprot datasets for identifying hydroxyproline sites.

| Metrics values for dbptm dataset |  |  |  |  | Metrics values for uniprot dataset |  |  |  |
| --- | --- | --- | --- | --- | --- | --- | --- | --- |
| Tests | Sn (%) | Sp (%) | Acc (%) | MCC | Sn (%) | Sp (%) | Acc (%) | MCC |
| Self – Consistency | 99.30 | 98.20 | 99.31 | 0.96 | 99.48 | 98.84 | 98.69 | 0.95 |
| Cross – Validation | 98.95 | 95.87 | 97.85 | 0.94 | 94.87 | 95.20 | 95.06 | 0.92 |
| Jackknife | 98.90 | 95.82 | 97.80 | 0.91 | 94.82 | 95.15 | 95.01 | 0.90 |

### Comparison Analysis

Observe Table 2 for a comparison analysis with the existing techniques “iHyd-PseAAC” [8], and “iHyd-PseCp” [25]. The comparison was also made with the most recent publication “iHyd-PseAAC (EPSV)” [32] for identifying the hydroxyproline sites. All these techniques attained the metrics records, employing the jackknife test method. It can be noticed from Table 2 that the accuracy (Acc), stability (MCC), sensitivity (Sn), and specificity (Sp) assess measured via operating proposed scheme are more predominant than the those values given by the former methodologies. There are two benchmark datasets borrowed from (a) dbptm and (b) uniprot database for comparison purpose with the existing schemes. Indeed, the newly proposed methodology is absolutely better suggestion throughout the past methodologies. There are a number of scientific and theoretical reasons can be explained for the improved quality of the developed scheme. Few of them are covered here. First of all, the proposed formulation is established by incorporating position and composition of primary protein structure and is beneficial to deal with the different length protein sequences in a thoughtful manner without missing any hidden data and also organize pairwise couplings in every possible permutation of amino acid residues. Second, it produces uniform dimension vectors, which contribute invariant size feature vectors that uniformly classify proteins corresponding to their properties. This concept allows the predictor to meticulously separate and appropriately distinguish each instance. Third, the correlation aspect is the principle concept that impart for computing feature vector. It has been assembled by considering each attribute group. Each expression deals with some specific metric and statistical measures. For the sake of convenience, every property of amino acids was standardized numerically within a suitable range. Also, it has been noticed that in comparison with previous methods proposed, the predicted outcomes are more superior and better than the former prediction rate.

**Table 2.** A comparison analysis of the proposed predictor with the existing predictors using well-known jackknife validation tests for the metrics given in Eq. (17). A comparison is made for jackknife test using benchmark datasets obtained from (a)dbptm and (b)uniprot database sources. It can be seen that the results obtained by using proposed predictor “iHyd-ProSite” is much better than all previous methodologies.

| Comparison Table |  |  |  |  |
| --- | --- | --- | --- | --- |
| *Predictors* | Sn (%) | Sp (%) | Acc (%) | MCC |
| *iHyd – PseAAC* | 80.66 | 80.54 | 80.57 | 0.51 |
| *iHyd – PseCp* | 86.35 | 99.12 | 96.58 | 0.89 |
| *iHyd – PseACC(EPSV)<sup>a</sup>* | 98.68 | 94.82 | 96.80 | 0.90 |
| *iHyd – PseACC(EPSV)<sup>b</sup>* | 97.02 | 94.57 | 96.01 | 0.88 |
| *iHyd – ProSite<sup>a</sup>* | 98.90 | 95.82 | 97.80 | 0.91 |
| *iHyd – ProSite<sup>b</sup>* | 94.82 | 95.15 | 95.01 | 0.90 |

## Discussion

Table 2 explain that, the values of sensitivity, specificity, accuracy and methew correlation coefficient for proposed predictor are higher than all the values obtained by utilizing former schemes. Sensitivity test describes the correctly predicted hydroxylated sites which are extraordinary larger than all reported values for previous methodologies. Also the stability of the predictor is measured by MCC value, and it can be observed that MCC values obtained by using proposed scheme are greater than above reported values. Undoubtedly, the proposed scheme is much helpful for diagnosing the biological problems efficiently.

## Conclusion

The novel proposed technique “iHyd-ProSite” is a new predictor to find hydroxyproline sites in protein sequences. Undoubtedly, it can be observed from the comparison Table 2 that the results obtained by using the proposed method are higher-up than all previous methods. For example, the accuracy calculated with the proposed tool (iHyd-ProSite) were 97.80 and 95.01 corresponding to two benchmark datasets obtained from databases (a) dbptm and (b) uniprot which is superior than the accuracies obtained by all previous predictors. Also MCC value were 0.91 and 0.90 with is greater than all schemes iHyd-PseAAC, iHyd-PseCp and iHyd-PseAAC (EPSV). Also the set of two data sets taken from dbptm and uniprot database were utilized for the proposed predictor validation. This technique is convenient to handle all types of biological data and can gently classify the unpredictable biological sequences. If the researchers are interested in the classification problems they should use this handy predictor, it can be helpful for future prediction problems.

### Bold the title sentence

Add descriptive text after the title of the item (optional).

## Conflict of Interest

The authors declare no conflict of interest, financial or otherwise.

